# Genome characterization, prevalence and transmission mode of a novel picornavirus associated with the Threespine Stickleback (*Gasterosteus aculeatus*)

**DOI:** 10.1101/502195

**Authors:** Megan A Hahn, Nolwenn M Dheilly

## Abstract

The complete genome sequence of an RNA virus was assembled from RNA sequencing of virus particles purified from threespine stickleback intestine samples. This new virus is most closely related to the Eel Picornavirus and can be assigned to the genus *Potamipivirus* in the family Picornoviridae. Its unique genetic properties are sufficient to establish a new species, dubbed the Threespine Stickleback Picornavirus (TSPV). Due to their broad geographic distribution throughout the northern hemisphere and parallel adaptation to freshwater, threespine sticklebacks have become a model in evolutionary ecology. Further analysis using diagnostic PCRs revealed that TSPV is highly prevalent in both anadromous and freshwater populations of threespine sticklebacks and is transmitted vertically to offspring obtained from *in vitro* fertilization in laboratory settings. It is thus necessary to test the impact of TSPV on the biology of threespine sticklebacks as this widespread virus could interfere with the behavioral, physiological, or immunological studies that employ threespine sticklebacks as model system.

**Abstract Importance:** The threespine stickleback species complex is an important model system in ecological and evolutionary studies because of the large number of isolated divergent populations that are experimentally tractable. For similar reasons, its co-evolution with the cestode parasite *Schistocephalus solidus*, its interaction with gut microbes, and the evolution of its immune system are of growing interest. Herein we describe the discovery of a RNA virus that infects both freshwater and anadromous populations of sticklebacks. We also show that the virus is transmitted vertically from parent to offspring in laboratory settings, suggesting that most experiments using sticklebacks from these lakes, and potentially others, were conducted in presence of the virus. This discovery can serve as a reminder that the presence of viruses in wild-caught animals for laboratory purpose is possible, even when animals appear healthy. Regarding threespine sticklebacks, the impact of TSPV on the fish biology should be investigated further to ensure that it does not interfere with experimental results.

## Introduction

The family Picornaviridae encompasses positive strand RNA viruses whose genome encodes a single large protein precursor (polyprotein). The typical genome organization comprises 6,700 to 10,100 nt and is composed of a 5’-untranslated region (UTR) that possesses an internal ribosome entry site (IRES) recognized by host cell ribosomes and allowing cap-independent translation, a single open reading frame encoding the polyprotein, and a 3’-UTR with a polyadenylated tract of variable length. The typical polyprotein is composed of facultative non-structural Leader (L^pro^), followed by a P1 region encoding three structural proteins (1AB that is cleaved into VP4 and VP3 in most picornaviruses, and 1C and 1D that encode for proteins VP2 and VP1 respectively), and two non-structural regions P2 and P3. Region P2 encodes proteins 2A^pro^ and 2B that impair the transport of host proteins, and 2C^ATPase^, a multifunctional ATPase involved in vesicle formation. Region P3 encodes proteins 3A that mediate presentation of membrane proteins during viral replication, 3B^VpG^, viral protein genome-linked that acts as a primer for RNA synthesis during RNA replication, 3C^pro^, a cysteine protease (picornain) responsible for maturation cleavage of the precursor polyprotein, and 3D^pol^, the RNA-dependent RNA polymerase. The gene regions coding for the capsid proteins and the non-structural proteins 2C, 3C^pro^ and 3D^pol^ are present in all described picornaviruses whereas the other non-structural proteins L, 2A, 2B. 3A and 3B^VpG^ may be absent in some species [1].

The family Picornaviridae is comprised of 40 genera and contains more than 94 species (as of February 2018), but many of these viruses are currently awaiting classification (http://www.picornaviridae.com). Until recently, only six species of picornaviruses had been reported in fish; the Eel Picornavirus 1 (EPV-1) in *Anguilla anguilla* [2], the Carp Picornavirus 1 (CPV-1) and Fisavirus (partial sequence) in *Cyprinus carpio* [3, 4], the Bluegill Picornavirus (BGPV-1) in *Lepomis macrochirus* [5], the Fathead Minnow Picornavirus (FHMPV-1) in *Pimephales promelas* [6], and a partial sequence of the Tioga Picorna-like virus in *Salvelinus fontinalis* [7]. The BGPV, CPV and FHMPV have recently been classified in the new genus *Limnipivirus*, whereas the EPV is the only member of the new genus *Potamipivirus*. A recent investigation of vertebrate RNA viruses led to the discovery of an additional 27 picornaviruses from diverse fish species [8]. The low similarity between the conserved precursor proteins and other picornaviruses suggest that within each fish species, a novel species of picornavirus was discovered.

The threespine stickleback, *Gasterosteus aculeatus* (hereafter ‘stickleback’) is a small teleost fish found widely distributed across all continents in the Northern hemisphere [9]. Sticklebacks have both anadromous and freshwater representatives, the latter of which have had recurrent adaptation in multiple different lakes [9, 10]. Freshwater sticklebacks are of particular interest in ecological and evolutionary studies because each lake comprises an independent population of fish that display population specific traits within only a few generations [11]. In particular, the region surrounding Anchorage, Alaska is of great interest because of the large number of lakes and distinct lineages of sticklebacks. Indeed, over the past 20 years this region has become a ‘hotspot’ for evolutionary research and for studies of host-parasite coevolution [12–20]. However, information on stickleback viruses remains scarce. The presence, and prevalence of DNA viruses has been scarcely investigated while the presence of RNA viruses remains unknown [21–23].

Here, we investigated the presence of viruses in the gut of apparently healthy sticklebacks. Using a combination of Illumina sequencing and complementary PCR we sequenced the full-length genome of a positive strand RNA virus. The predicted polyprotein displays the typical organization of a picornavirus and genome sequence composition and phylogenetic analyses suggest that it belongs to the same genus as EPV. The virus was provisionally named Threespine Stickleback Picornavirus (TSPV). Additional targeted PCRs were used to test prevalence in marine and freshwater populations of sticklebacks from Alaska and to test the potential for vertical transmission of TSPV to offspring.

## Materials and Methods

### Virus isolation

Viruses were initially purified from the intestine of five apparently healthy *G. aculeatus* collected in Cheney Lake, Alaska (61° 12′ 17″ N, 149° 45′ 33″) in June 2016. Viruses were purified using a filter-chloroform nuclease virus purification protocol. Briefly, tissue samples were homogenized in sterile PBS by bead beating with 3 mm glass beads. The homogenates were centrifuged for 1 min at 6000 rpm. The pellets were discarded, and the solutions were further diluted with 500 µl of PBS and centrifuged at 6000 rpm for 6 min to remove remaining cell debris. The supernatants were then filtered successively through a 0.4 µm filter and 0.22 µm filter and incubated for 10 min in 0.2 volumes of chloroform. The viruses were then separated from the chloroform by centrifugation for 20 seconds at 20,000 rpm. A second chloroform treatment was applied to ensure removal of bacterial contaminants. The purified viruses were finally treated with 2.5U of DNase I and 0.25 U of RNase at 37°C for 3 hours to eliminate non-viral DNA and RNA. EDTA was added at a final concentration of 20mM.

### Sample preparation, sequencing and assembly

Initially, both viral DNA and viral RNA were extracted using the QIAamp Mini Elute Virus Spin Kit according to manufacturer’s instructions. DNA was then removed using a Turbo DNAse treatment. The True Seq mRNA library preparation kit was used before single-end sequencing, for 100 cycles, on an Illumina Hi-Seq 4000 (Institute of Biotechnology at Cornell University). For each dataset, adapters were removed using Trimmomatic and PhiX174 contaminants were removed using Bowtie 2 (--very-sensitive-local). We obtained 2.38, 4.32, 6.98, 8.18 and 9.70 million high quality reads for each respective sample. Then, for each sample, d*e novo* assembly was completed using Trinity [24]. Partial sequences of a virus that showed high similarity to EPV were assembled independently from all five samples. Reads from the five samples were thus pooled to improve viral *de novo* assembly. Eight contigs of more than 500 nt showed high similarity to EPV. The two longest contigs had 6741 nt and 1536 nt and were assembled in a single draft genome of 8267 nt nucleotides, with potentially missing nucleotides at the 5’ and 3’ ends.

### Fragment amplification and Sanger sequencing

First strand cDNA was synthesized by reverse transcribing 500 ng of RNA from purified viruses and prepared with 0.2 µg/µl Random Hexamer Primer in a 20µl reaction volume containing 40 U/µl Ribolock™ RNase Inhibitor, 1 mM dNTPs, 200 U/µl RevertAid H Minus Reverse Transcriptase (Thermo Fisher Scientific, Waltham, MA, USA), and water, as per manufacturers recommendations. Polymerase chain reaction was conducted using the Advantage 2 PCR system (Invitrogen) and primers from table 1. To complete the 3′ ends of the genome sequence, we performed a Rapid Amplification for cDNA End (RACE) using Invitrogen 3’RACE Systems following the manufacturer’s instructions. PCR products were sequenced using Sanger sequencing.

### Phylogenetic analyses

To investigate the relationship of TSPV with other related picornaviruses, TSPV protein sequences from the P1, 2C and 3CD regions were aligned using clustalo with representatives from each genus, and with all known picornaviruses of fishes. Maximum likelihood phylogenetic trees were constructed in Seaview4 [25] using the LG substitution model [26] implemented in PhyML [27] with the nonparametric Shimodaira-Hasegawa-like procedure. Pair-wise identity matrices were obtained using Clustal-Omega online (https://www.ebi.ac.uk).

### Sampling, sample preparation and diagnostic PCR

In June of 2018, we collected intestines of sticklebacks from an anadromous marine population in Rabbit Slough (61° 32′ 08.1″ N 149° 15′ 10.0″ W), and from isolated freshwater populations from Cheney Lake, Wolf Lake (61° 38′ 36″ N, 149° 16′ 32″ W), and Loberg Lake (61° 33′ 33.5″ N 149° 15′ 28.9″ W). Upon dissection, intestines were immediately transferred in RNA later and preserved at –80 °C until use. Total RNA was extracted using Trizol following the recommended protocol. We also collected mature males and gravid females from Rabbit slough, Cheney Lake, and Loberg lake and completed crosses *in vitro* to obtain families. Eggs were washed in methylene blue and maintained in individual Petri dishes with daily water changes until hatching. Following hatching, six one day old larvae from each lake were collected and preserved in RNA Later. Total RNA was extracted using the RNeasy kit (CAT# 74104) following the manufacturer’s guidelines.

Diagnostic PCRs were completed as described above using primers targeting the conserved RNA-dependent RNA polymerase gene (F, 5’ TCT CCT ACC AAA CCC GCA AC 3’; R, 5’ TTC CTC CGC CAC CAG ATA GA 3’). Amplicon presence was assayed with 1% agarose gel with SyBR Safe.

### Ethics statement

Stickleback collection followed guidelines for scientific fish collection by the State of Alaska Department of Fish and Game in accordance with Fish sampling permit #P17-025 and #P-18-008 and fish transport permits 17A-0024 provided to NMD. Fish were maintained at Stony Brook University under the License to collect or possess #1949 provided by the New York State Department of Environmental Conservation to NMD. Fish experiments were conduct following protocols described in Institutional Animal Care and Use Committee (IACUC) #237429 and # 815164 to Michael Bell and NMD, respectively. Fish euthanasia was conducted using MS222 and decapitation before parasite and tissue sampling. All experiments were performed in accordance with relevant guidelines and regulations in the Public Health Service Policy (PHS) on Humane Care and Use of Laboratory Animals.

## Results and discussion

### Molecular characteristics of TSPV

The genome organization of TSPV-1 is depicted in Figure 1 and can be described as follows: VPg+5’UTR[1AB-1C-1D-2A1^NPGP^/2A2^NPGP^/2A3^H^-Box/NC-2B-2C^Hel^/3A-3B^VPg^-3C^pro-^3D^pol^]3’UTR-poly(A)

**Figure 1:**
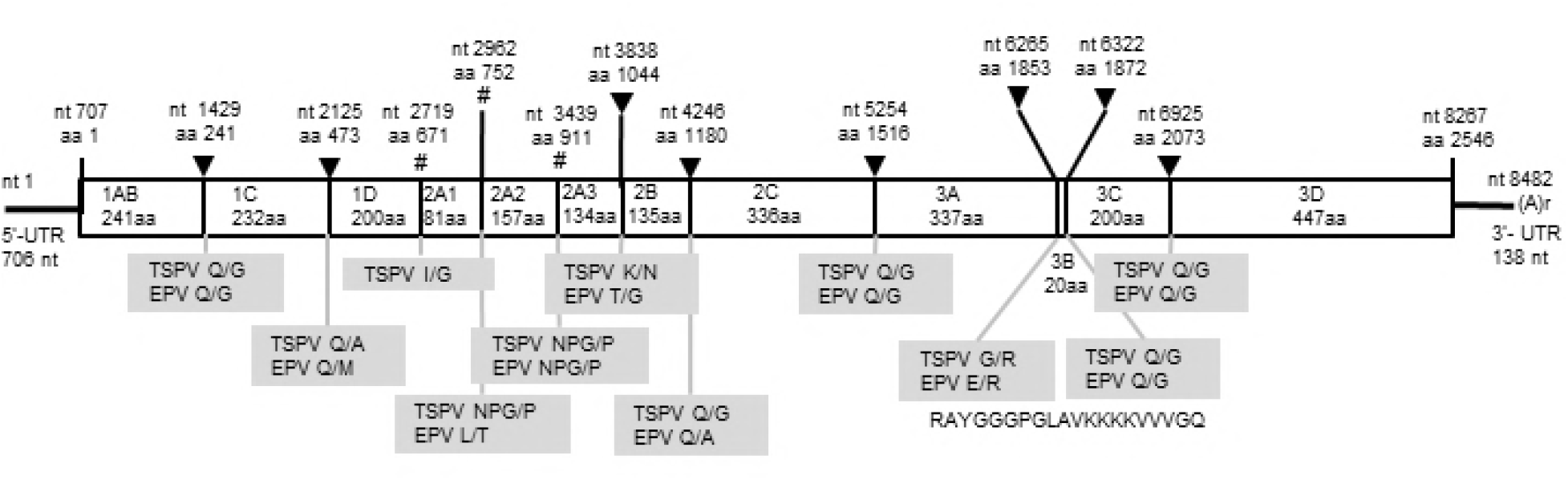
Schematic diagram of the predicted TSPV genome structure. Numbers on the top indicate amino acid (aa) position and nucleotide (nt) positions. The 5’ and 3’ UTRs (line) and the open reading frame (box) are displayed. Predicted protease cleavage sites are indicated with triangles (▾) and ribosomal skip sequences are indicated with the hash (#). Grey boxes show a comparison of the cleavage sites with EPV-1.

The positive strand RNA genome is composed of a 5’-UTR of 707 nt, an ORF of 7560 nt, a 3’-UTR of 138 nt and a poly A tail of at least 50 nt and is overall more similar to EPV than to any other picornavirus (table 1).

**Table 1:**
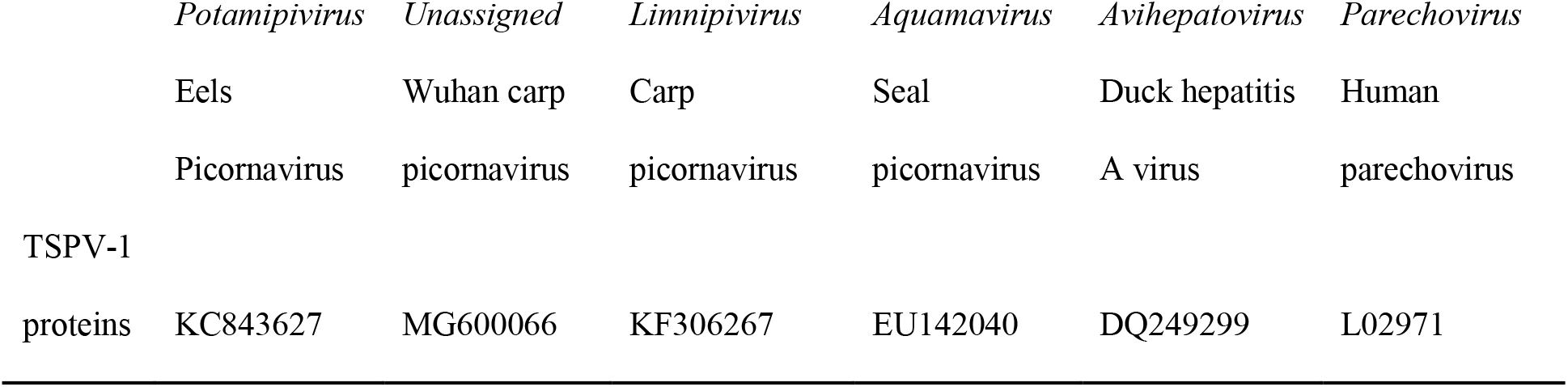
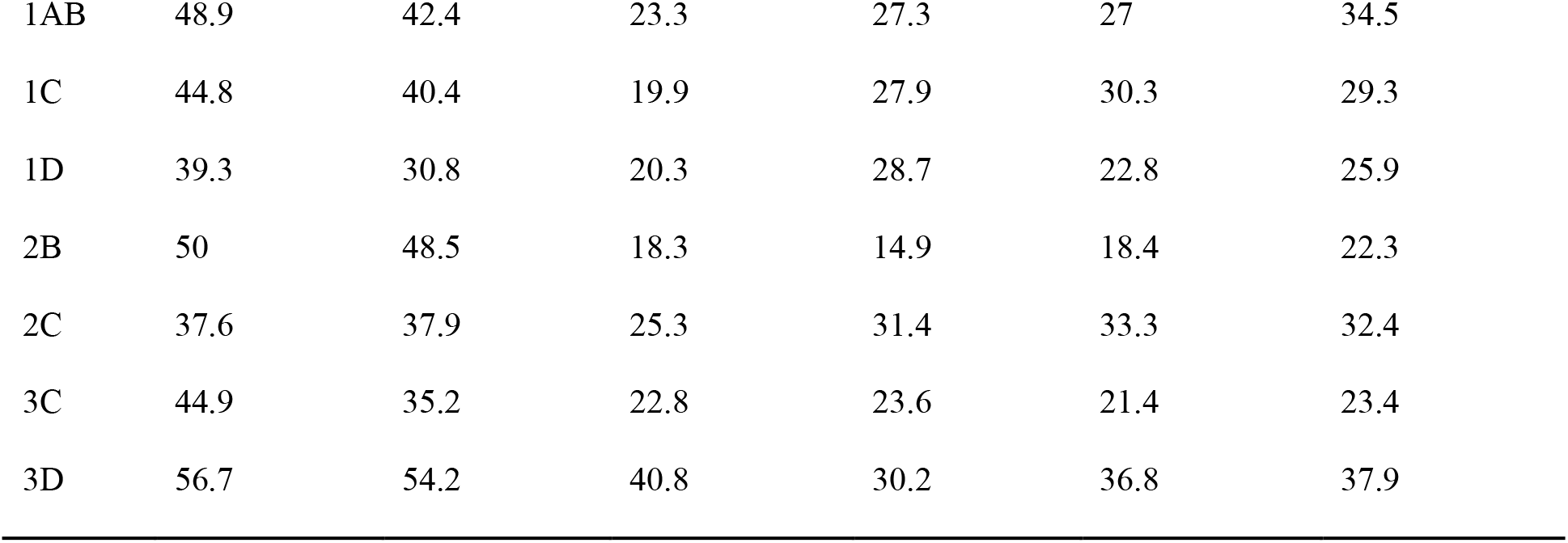
Percent pairwise amino acid identity of TSPV proteins to other picornavirus proteins from the genus *Potamipivirus, Limnipivirus, Aquamavirus, Avihepatovirus*, and *Parechovirus*.

The deduced polyprotein precursor was 2520 aa long with an overall base composition of 28.6% (A), 19.2% (C), 25.4% (G), and 26.8% (T). The genome of TSPV had no obvious leader sequence. It had three capsids (1AB, 1C and 1D). Conserved rhv_like hydrophobic domain of capsid (cd00205) proteins were found within proteins 1AB and 1C (aa positions 103–228 and 294–465 respectively).

We identified five non-structural proteins (2A1, 2A2, 2A3, 2B, and 2C) within the P2 region. The 2A region is complex with high similarities with EPV, CPV-1 and BGPV-1 (Figure 2). Similarly to CPV-1 and BGPV-1, TSPV encodes two 2A proteins that end with an NPG/P ribosome skipping motif that is found in many Picornaviruses and mediates in *cis* co-translational termination–re-initiation of RNA translation. The 2A3 gene region exhibits a parechovirus-like and Avihepatovirus-like structure including the conserved H-box and NC-box. The 2B protein is also homologous to parechovirus and avihepatovirus. A conserved RNA helicase domain was found within protein 2C (aa positions 1316–1422, pfam00910) that contained the three domains originally identified by Koonin and Dolja [28].

**Figure 2:**
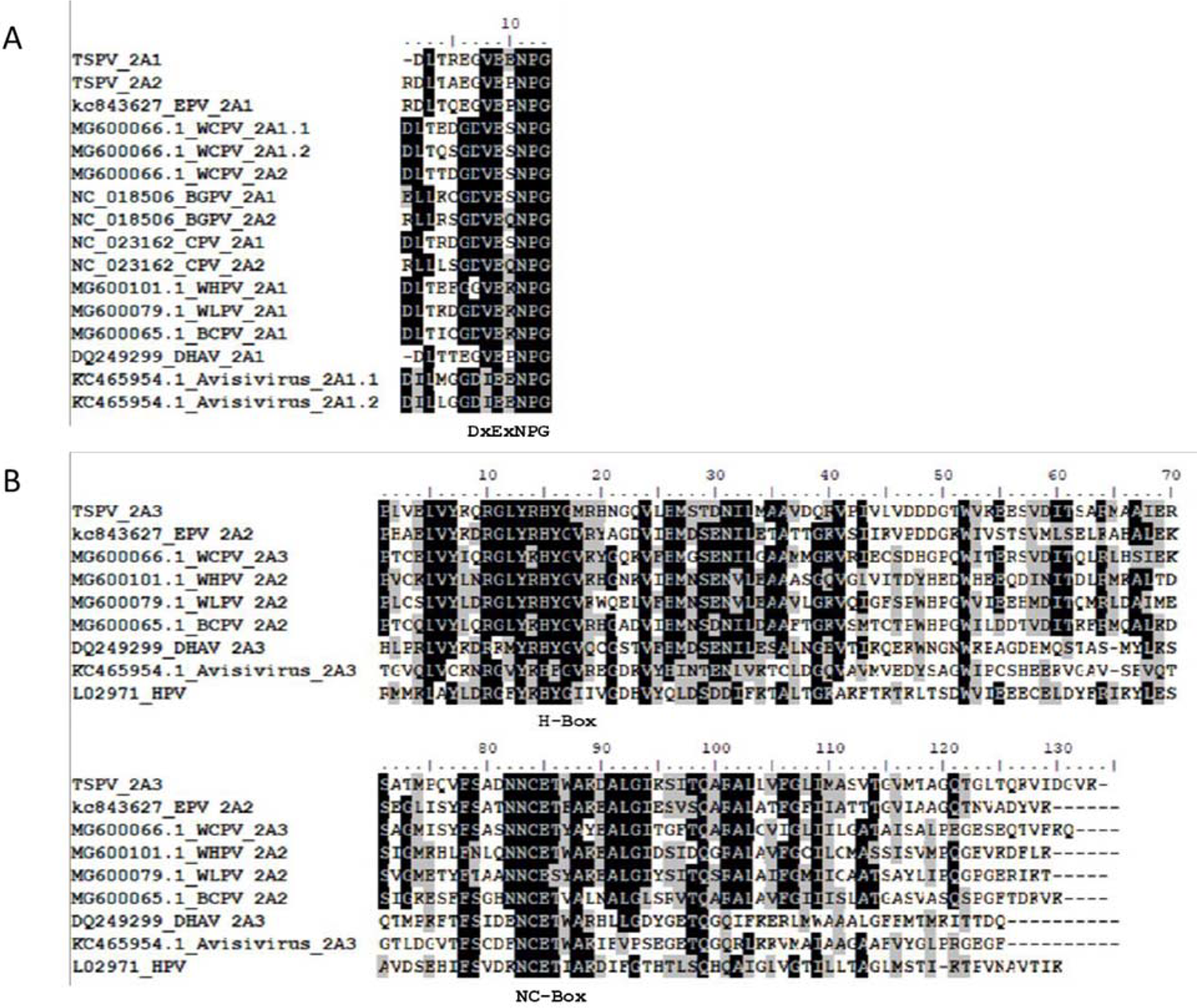
sequence alignment of TSPV 2A proteins with proteins 2A from related species. A/ Alignment of the Aphtovirus-like 2A1 and 2A2. B/ Alignment of the Parechovirus-like 2A3. Characteristic functional motifs are indicated.

The P3 region contained three non-structural proteins (3A, 3B^VpG^, 3C^pro^ and 3D^pol^). 3A showed no obvious homology and was twice as long as 3A in related picornaviruses. It showed no homology to the EPV 3A region either. The N terminus of 3B exhibits the characteristic RAY of EPV, CPV-1, BGPV-1 and Parechoviruses. A conserved 3C cysteine protease was found in protein 3C (aa positions 1954-2056, pfam00548). The putative protease has the core domain of picornaviruses (GxCG) and the GxHxxG substrate binding pocket. A conserved RdRP domain was found in protein 3D (aa positions 2089–2487, cd01699, pfam00680). The RdRp had all eight motifs of the core region described by Koonin and Dolja [28, 29].

### Phylogenetic analyses

To investigate the relationship of TSPV with EPV and other picornaviruses, protein sequences from the P1 and 3CD regions were aligned with representatives from each described genus and additional unassigned candidate picornavirus species from fish (Figure 4). Maximum likelihood phylogenetic trees confirmed that TSPV was most closely related to EPV and belongs to the genus *Potamipivirus*. It is the first known RNA virus of threespine sticklebacks.

**Figure 4:**
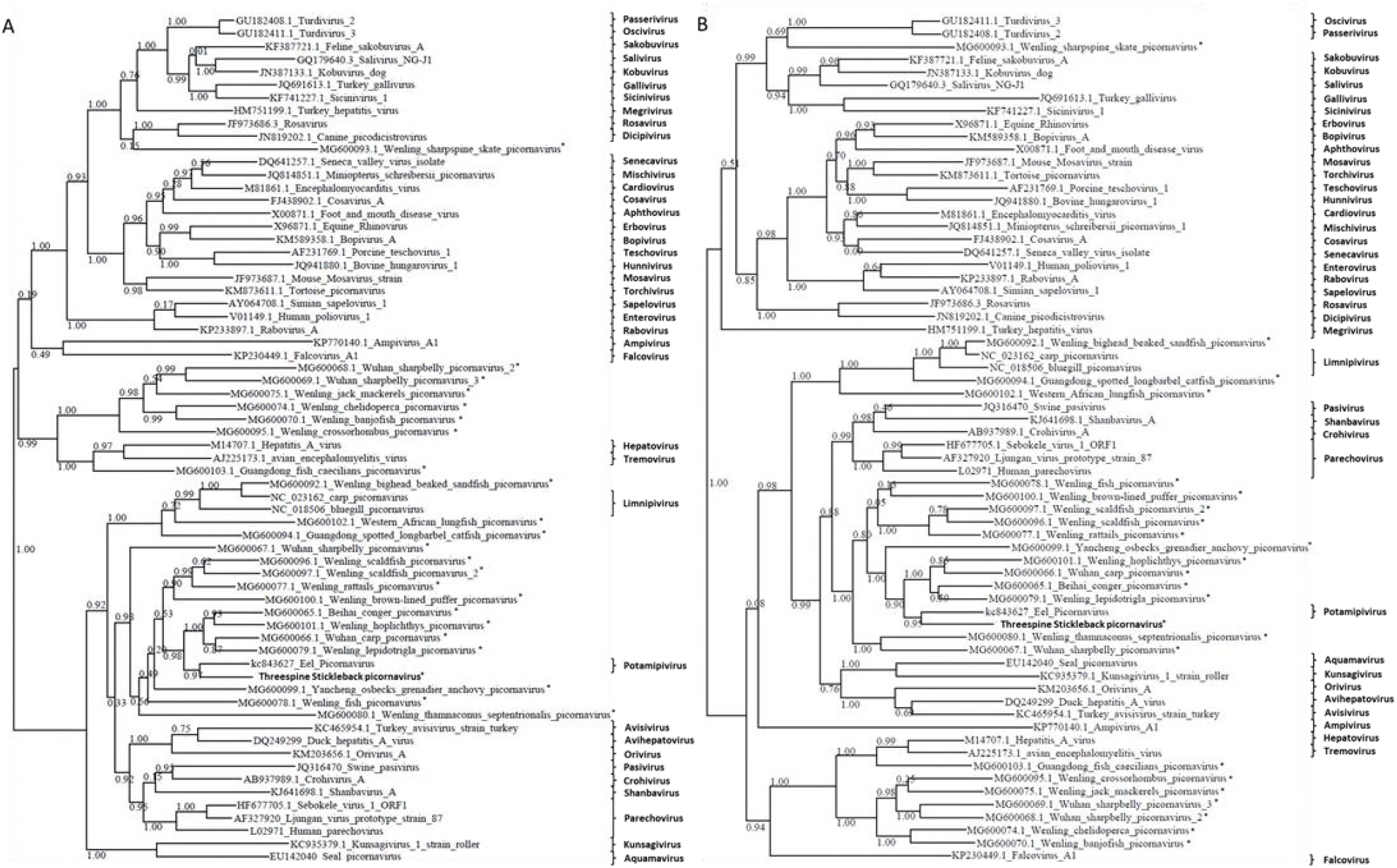
Phylogenetic analysis of picornavirus 3CD (A) and P1 (B) gene regions. One representative sequence of each of the 40 described genera, and an additional 22 sequences of fish-associated viruses obtained from GenBank and the TSPV were included. The trees were inferred with PhyML using the LG substitution model. Values next to the branch indicate the result of a Shimodaira-Hasegawa branch test.

### Distribution and transmission

Diagnostic PCR on RNA extracted from intestines of sticklebacks from anadromous and freshwater populations in Alaska revealed high prevalence of TSPV: 100% prevalence was observed in anadromous fish collected from Rabbit Slough (20 individuals) and in freshwater fish from Cheney lake (32 individuals) and Loberg Lake (15 individuals) and 88% in freshwater fish from Wolf lake (17 individuals).

We used *in vitro* fertilization to cross males and females from Rabbit Slough, Cheney Lake, and Loberg lake. Eggs were washed in methylene and maintained in individual Petri dishes with daily water change until hatching. We then tested TSPV presence in one day old larvae from six families for each population and found TSPV in all samples indicating that the virus is vertically transmitted from parents to offspring obtained in laboratory conditions.

Vertical transmission of picornaviruses has previously been observed in both invertebrates and vertebrates [30–35]. While picornavirus-like particles have been found in carp, bluegill, baitfish sea bass, barramundi, smelt, turbot, and salmonids, vertical transmission has not been routinely been tested [4, 5, 36–38]. However, picornavirus-like particles have been found in the ovarian fluids of cutthroat, brown, and brook trout suggesting vertical transmission [38]. Given the high prevalence of TSPV in our field caught samples and demonstration of vertical transmission in laboratory crosses, it is very likely that lab raised individuals from these populations commonly carry the virus.

Viruses in fish have most commonly been investigated when mortality is occurring in aquaculture stocks or fisheries, but it is also necessary to investigate the presence of viruses in healthy fish to better reconstruct virus phylogenies and understand processes of pathogenicity [39]. Regarding the use of threespine sticklebacks and other fish species in laboratory settings as model systems in evolutionary ecology and immunology, it is crucial to monitor virus presence and investigate the impact of TSPV and other highly prevalent viruses on fish behavior, physiology and immunity.

### Data availability

The TSPV genome sequence was submitted to Genbank under the Nucleotide Accession number MK189163 and sequencing data were submitted to NCBI Sequence Read Archives under Bioproject accession numbers SAMN10346744, SAMN10346743, SAMN10346742, SAMN10346741 and SAMN10346745.

## Acknowledgments

We would like to acknowledge the expertise and assistance of Dr. Michael Bell in fish collection and dissection techniques. This project was supported by the Eppley Foundation for Research and the Maze-Landeau Foundation.

